# Tunable porosity in a hydrogel with extreme vibration damping properties

**DOI:** 10.1101/2025.01.16.633355

**Authors:** Graham J. Day, Qicheng Zhang, Chrystel D. L. Remillat, Gianni Comandini, Adam W. Perriman, Fabrizio Scarpa

**Affiliations:** School of Cellular and Molecular Medicine, University of Bristol, BS8 1TD Bristol, UK; Bristol Composites Institute, School of Civil, Aerospace and Design Engineering (CADE), University of Bristol, Bristol BS8 1TR, UK; Centre for the Cellular Microenvironment, Division of Biomedical Engineering, James Watt School of Engineering, The Advance Research Centre, University of Glasgow, Glasgow G12 8QQ, UK; Research School of Chemistry and John Curtin School of Medical Research, Australian National University, Canberra ACT2601, Australia

**Keywords:** hydrogel, vibration, mechanics, alginate, poloxamer, dynamic properties

## Abstract

We have developed hydrogel systems with tunable porosity through dialysis casting by varying alginate and poloxamer compositions from 0% to 10%. These gels feature diverse porosity topologies, yielding loss factors between 16% and 29% in the 50 Hz to 300 Hz frequency range. The dynamic modulus shows a remarkable increase of over an order of magnitude, reaching approximately 3 MPa compared to the static modulus. Vibration transmissibility tests and dynamic mechanical analysis reveal that the poroelastic and pneumatic-like effects from the tunable porous structures contribute significantly to this damping effect. Furthermore, these hydrogels are biosourced and biodegradable, providing a sustainable alternative to conventional fossil-based damping materials.

## Introduction

Hydrogel systems are materials extensively used in the cosmetic industry, biomedical and tissue scaffolding applications^1^. The viscoelastic properties of hydrogels are extensively evaluated because of their importance in rheology and bioprinting applications. The damping capacity of hydrogel systems is also cited as a proof of their energy absorption properties under large quasi-static and dynamic loadings^2, 3^. Vibrations are a subset of dynamic loads that are critical to the structural integrity of machinery, airframe and structures associated to transport applications and constructions. Vibration damping technologies have been developed during the last eight decades for the suppression, alleviation and control of loads and deformations generated by ambient or forced vibrations^4, 5^. Damping technologies make use of the synergy between the dissipative and viscoelastic properties of the constituent materials and the substrates/packages that support those materials systems. Most materials traditionally used in damping technologies are of fossil origin, and include thermoset and thermoplastic polymers, as well as elastomers/rubbers. Biobased materials offer however an alternative approach in designing devices for vibration damping technologies, as well as providing enhanced sustainability and reduction of the environmental impact of those applications. Hydrogels, in particular, can be produced using biobased and biodegradable compounds and offer very interesting performance in terms of damping capacity. Du et al have observed recoverable energy dissipation in drop-tower tests carried out on sodium alginate-silicon nitride PVA composite biomaterials subjected to ∼ 3J and ∼6 J of kinetic energy^6^. Polydimethylsiloxane gels (PDMS) have been used to fill polymeric and piezoelectric porous skeleton and generate three-dimensional composites with significant damping capabilities. PZT-PDMS composites have shown loss factors of ∼20% within 1–20 Hz sweeps performed with dynamic mechanical analyzers (DMA)^7^. Polyurethane-carbon nanotube-PDMS gel composites have also exhibited loss factors over 24% during vibration transmissibility tests within 50–600 Hz^8^. The characterization of hydrogel materials from a purely vibration damping perspective has received however some limited attention. Wang et al have analyzed polyacrylamide (PAAm) and PDMS gels using a bending rig with base acceleration excitation. The viscous damping ratios of the PAAm gels associated to the first bending mode were between ∼2% and 3%, while those of the PDMS beams were close to 8%^2^.

Alginate-Poloxamer systems have shown some promise as bioink platforms for tissue engineering and scaffolding, also in view of their bioprinting capabilities^9^. The multiscale porosity of those hydrogel systems can be tailored with the poloxamer concentration, leading to different mechanical performance and poroelastic effects. Poroelasticity also affects the vibration damping properties of porous materials^10^, but also impacts the transport properties of the solvent through the polymeric network of the gels, contributing to energy dissipation^2, 11^. In this work we show how alginate/poloxamer hydrogel systems can dramatically increase their dynamic stiffness when subjected to vibration loading, up to values close to 3 MPa and loss factors ∼ 0.28 within the 100 Hz – 300 Hz range.

## Materials and Methods

### Production of the hydrogels

Pre-gel solutions with a mass of 50 g were produced by adding 25 g of ultra-pure water to a PP30-60ML Mix Cup (Hauschild-SpeedMixer, Germany), then adding the calculated masses of dry sodium alginate and Poloxamer 407 powders (**Supplementary table S1**). The powders were passed through a sieve as they were added to remove clumps. Finally, the mixture was topped up to 50 g with ultra-pure water and sealed with the lid. The solution was mixed using a dual-asymmetric centrifuge-150 (DAC150, Hauschild-SpeedMixer, Germany) at 250 rpm for 5 mins at room temperature. The vessels were then left to stand on the benchtop for 15 mins to give the powders time to hydrate. Finally, the pre-gel solution was mixed in the DAC150 at 2500 rpm for 10 mins to fully homogenize.

A dialysis mold was inserted into a wetted dialysis tubing cellulose membrane (MWCO 14000), with a width of 43 mm (Scientific Laboratory Supplies, UK) and clipped at the bottom of the mold. The pre-gel was then poured into the dialysis tubing to cover the dialysis mold and pockets of air were removed by manipulating the membrane. The tubing was then sealed by clipping at the top of the dialysis mold. The loaded membrane was then placed into 1.5 L of pre-warmed 37 °C ultra-pure water (in a 2 L beaker) and incubated at 37 °C for 1 h to induce the sol-gel transition of the Poloxamer. To cross-link the solution and form the final hydrogel, the beaker was removed from the 37 °C incubator and the solution stirred at 200 rpm with a magnetic stir bar, taking care to avoid knocking the dialysis membrane. Subsequently, 147.01 g of calcium chloride (CaCl_2_) was added to the stirring solution followed by 0.5 L of pre-warmed 37 °C ultra-pure water, giving a final CaCl_2_ concentration of 500 mM, and left to stir at room temperature for 18 h.

The cross-linked hydrogel with the embedded mold were removed from the dialysis membrane and a sharp knife was used to cut away excess hydrogel from the mold and release the hydrogel sample in the desired shape. To wash out excess Poloxamer 407, the final hydrogels samples were submerged under 200 mL ultra-pure water for 24 h, whilst replacing the water hourly over the first four hours.

### Cryogenic scanning electron microscopy

Cryogenic scanning electron microscopy (cryo-SEM) of the 5 wt% alginate hydrogels was conducted at the Wolfson BioImaging Facility, University of Bristol. Scanning electron micrographs were captured on a FEI Quanta 200 field emission gun SEM (Thermofisher Scientific, USA), under high vacuum. To image the samples in a condition as close to the hydrated state as possible, punches were taken from the hydrogel samples and frozen by plunging in a liquid nitrogen slurry. Then, in an environment-controlled chamber built on to the SEM, the sample was fractured and the surface sublimated for three minutes. The surfaces were coated with carbon prior to imaging.

Pore diameters were manually measured using FIJI software. Statistical analysis of the pores was performed on 290 measurements taken from three micrographs per hydrogel composition.

### Quasi-static uniaxial compression testing

Quasi-static uniaxial compression tests were carried out using an FMS-500-L2 Force Measurement System (Starrett, Australia) fitted with a 10 N load cell. Cylindrical dialysis-casted hydrogel samples, with a diameter of 12 mm and height of 5 mm, were loaded onto the bottom platen and the top platen was lowered until it contacted the top surface of the hydrogel. The samples were then compressed at a rate of 1 mm·min^−1^ until the load cell recorded a force of 8 N, at which point it stopped compressing. The Young’s modulus (*E*) each of the hydrogels was determined from the linear portion of the stress–strain curve at 5–10% strain (**Equation 1**).

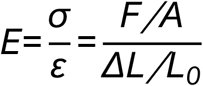

**Equation 1**. Determining the Young’s modulus, *E*, where *σ* is uniaxial compressive force (*F*; Newtons) per unit surface area (*A*; mm^2^) and *ε* is strain, or the change in height (*ΔL*; mm) divided by the original height (*L*_*0*_; mm).

### Vibration transmissibility tests

Hydrogel blocks of dimensions (*l×w×h*) 30×30×15 mm, were dialysis casted. The seismic vibration rig used for vibration transmissibility experiments has been previously described and characterized^10^, but will be described here. The rig consisted of a steel base plate mounted on to a vertically orientated electrodynamic shaker (LDS V406 Permanent Magnetic Shaker, LDS, UK), with the hydrogel samples mounted on the center of the base plate. The hydrogels were pre-strained by the placing a connecting plate on the top surface of the hydrogel, on to which additional masses (top mass) could be affixed with bolts. An adhesive agent (DEHP Alginate Tray Adhesive Liquid, Henry Schein, UK) was used to fix the hydrogel to the base plate and the connecting plate to the hydrogel. Two accelerometers (model 333 M07, PCB Piezotronics, USA) were screwed into the base plate and the top mass to measure the vibration signal before and after passing through the hydrogel. Holders, which could be attached on the sides of the base plate, were used to center the hydrogel, and facilitated the unscrewing and screwing of top mass plates while minimally disturbing the hydrogel. The five top masses used to pre-strain the hydrogels (including the masses of the connecting plate, accelerometer, and bolts) were 32.44 g, 64.25 g, 111.10 g, 160.43 g, and 209.13 g.

Digital white noise signals were generated by an in-house MATLAB code, which was converted to an analog signal by a NI USB-6211 data acquisition (DAQ) device (National Instruments, USA) and then amplified by a power amplifier (LDS PA100E) to drive seismic vibration of the base plate. The accelerometers measured the analog signals corresponding to vibration of the base plate (undamped, before passing through the hydrogel) or top mass (damped) and transmitted to a NI-9234 C Series Sound and Vibration Input module (National Instruments, USA), which converted the signal to digital. Finally, another in-house MATLAB code recorded the data.

The transfer function, TF(*ω*), dynamic modulus, *E*_*d*_, and dynamic loss factor, *η*_*d*_, were determined as previously described^10^. In brief, *η*_*d*_, was obtained from a simplified seismic base vibration model, where the sample is represented by a linear spring with stiffness (real value) *k*, mass *m*_*s*_ and a hysteretic loss factor η (**Equation 2**).

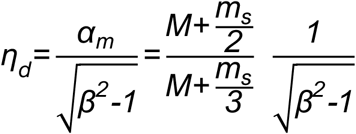

**Equation 2**. Calculating the dynamic loss modulus, *η*_*d*_, where *α*_*m*_ is the mass coefficient[ref], *β*^*2*^ is the peak amplitude of the transfer function at resonance and *M* is the inertia of the top mass.

Given that resonance frequency, *ω*_*n*_, is proportional to stiffness *k* (**Equation 3**)^10^, the dynamic storage modulus, *E*_*d*_, could be calculated at resonance (**Equation 4**). Accordingly, the dynamic loss modulus, *E*_*l*_, could be calculated from *η*_*d*_ and *E*_*d*_ (**Equation 5**).

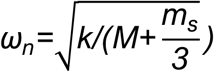

**Equation 3**. The relationship between resonance frequency, *ω*_*n*_, stiffness, *k*, top mass inertia, *M*, and sample mass, *m*_*s*_.

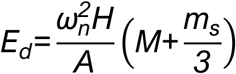

**Equation 4**. Determining the dynamic storage modulus, *E*_*d*_, at resonance frequency.

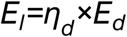

**Equation 5**. Determining the dynamic loss modulus, *E*_*l*_.

### Dynamic Mechanical Analyser

Dynamic mechanical analysis (DMA) tests were performed using a DMA 850 instrument (TA Instruments). Cylindrical hydrogel samples with a diameter of 12 mm and a height of 6 mm were tested under compressive loading. Frequency sweeps were conducted incrementally from 0.1 Hz to 200 Hz in overlapping ranges (e.g., 0.1–10 Hz, 9–20 Hz, etc.), with a new pristine sample from the same hydrogel batch used for each range to limit cumulative strain. A compression clamp with a diameter of 15 mm was utilized, and the test chamber temperature was maintained at a constant 25 °C. A conditioning preload force of 0.001 N was applied prior to testing. An oscillation amplitude of 10 μm was employed to ensure elastic sinusoidal compression of the samples.

## Results and discussion

### Hydrogel fabrication

A range of hydrogels with systematically increasing alginate (A) and Poloxamer 407 (P) compositions were produced to investigate the effect porosity had on the material stiffness. Increasing the alginate concentrations was expected to increase the material stiffness by increasing the network density and cross-linking concentration, whereas increasing the Poloxamer was expected to increase the pore size. Three alginate concentrations were chosen at 1.25, 2.5, and 5.0 wt% (A_1.25_, A_2.5_, and A_5_, respectively), with each group containing a 0.0, 2.5, 5.0, or 10.0 wt% Poloxamer (P_0_, P_2.5_, P_5_, and P_10_, respectively), giving 12 unique gel compositions. For example, the 5 wt% alginate with 2.5 wt.% Poloxamer hydrogel is hereby referred to as A_5_–P_2.5_. Calcium ions were chosen as the divalent metal ion necessary to cross-link the alginate chains (**Supplementary figure S1**). Hydrogel samples of defined and reproducible dimension were required for the compression and vibration transmissibility experiments, so were casted in specifically shaped moulds, however, initial attempts to cross-link with Ca^2+^ ions proved difficult, as the Ca^2+^ could not diffuse evenly through deep samples, resulting in uneven or incomplete cross-linking. To solve this issue, a casting method involving dialysis was conceived, involving 3D printed moulds that were open on two sides placed into a semi-permeable dialysis membrane, which would allow the Ca^2+^ to diffuse from both sides to halve the depth required for the ions to travel and reduce the effects of uneven cross-linking (**Figure 1**). Two types of moulds were designed: one to produce pucks (12 mm in height × 5 mm diameter) for compression testing (**Supplementary figure S2**); and blocks (30×30×15 mm) for vibration transmissibility experiments (**Supplementary figure S3**). The bars at the ends of the moulds indicated where the clips would be placed to seal the membrane, to ensure the volume of the membrane was constant across samples.

**Figure 1.**
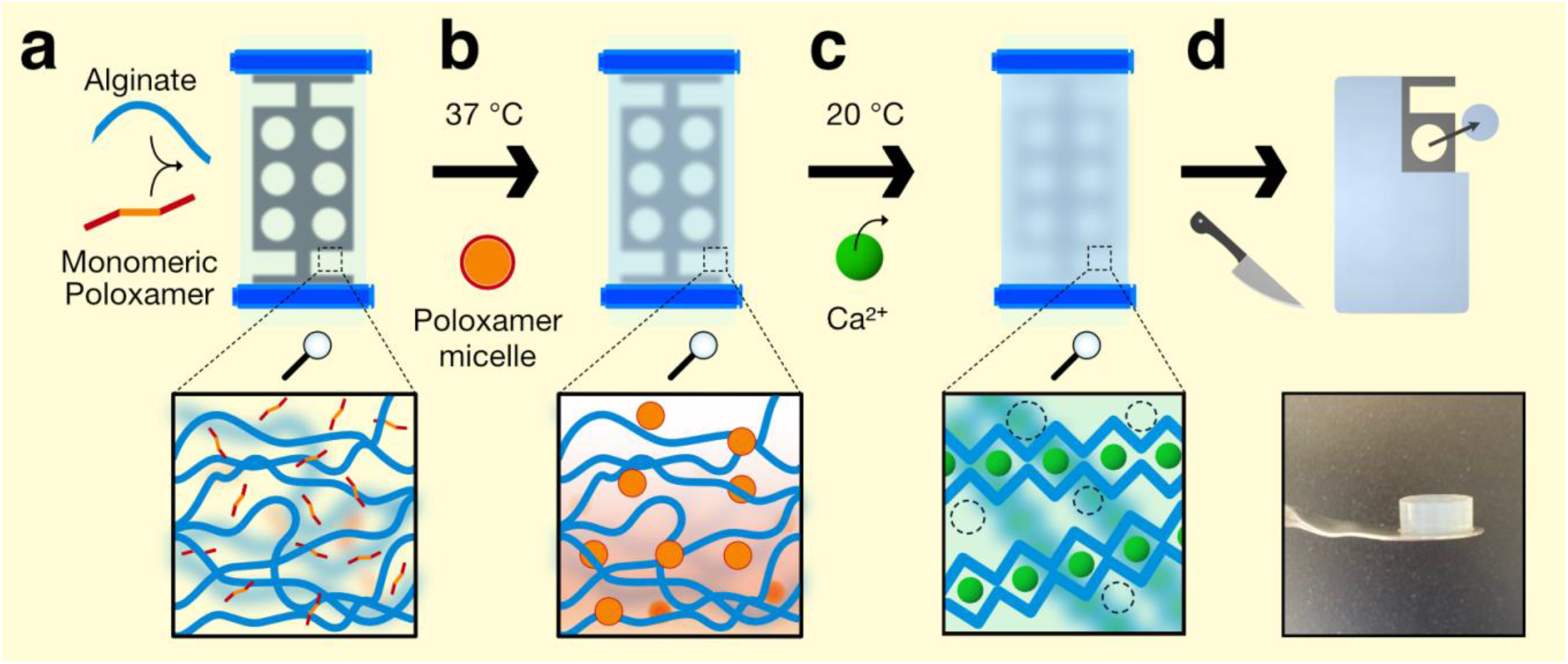
Schematic of dialysis-casting the hydrogels. **a** sodium alginate polymer (blue lines) was mixed with monomeric Poloxamer 407 (red–orange lines) in water at room temperature and added a dialysis tubing membrane (light blue) containing a plastic mould (dark grey), and then the dialysis tubing membrane was clipped shut (dark blue). **b** The membrane was submerged in water at 37 °C to induce a sol–gel transition of the Poloxamer, forming micelles (red–orange circles). **c** Calcium chloride was added to the water exterior to the dialysis membrane tubing to cross-link the alginate with Ca^2+^ ions (green circles). The solution was gradually cooled to 20 °C to reverse the sol– gel transition of the Poloxamer, so the Poloxamer could diffuse out of the membrane as Ca^2+^ diffused in. **d** Finally, after the alginate was fully cross-linked inside the dialysis membrane, excess hydrogel was cut away from the mould and samples of the desired dimensions were removed.

### Stiffness related to microstructure

The effect of the alginate and Poloxamer concentrations on the stiffness of the hydrogels was investigated *via* uniaxial quasi-static compression testing. The stiffnesses of the hydrogels increased with alginate concentration, with the measured Young’s modulus (*E*) being 21.9 ±2.4, 88.7 ±17.4, and 167.4 ±49.0 kPa for the A_1.25_, A_2.5_, and A_5_ hydrogels with no Poloxamer, respectively (**Figure 2b**, black bars). This stiffness increase was thought to arise from strengthening the alginate network with higher concentrations, leading to increased cross-linking density and thicker fibres^12^. For the low alginate A_1.25_ group, no significant variation in the stiffness of the hydrogels was observed with the addition of Poloxamer, although it is possible they phase separated during formation. For the A_2.5_ gels, *E* decreased from 100 to 10 kPa with increasing Poloxamer. The high alginate A_5_–P_0_, –P_2.5_, and P_5_ hydrogels were isoelastic with *E* = 167.4 ±49.1, 192.6 ±28.9, and 192.4 ±20.7 kPa, respectively. However, the stiffness of the A_5_–P_10_ was significantly lower at *E* = 64.8 ±19.4 kPa. Inspection of the stress–strain curves revealed the A_5_–P_0_, –P_2.5_, and P_5_ were linear up to the ∼30% strain measured and exhibited strain stiffening behavior (**Figure 2c i–iii**). In contrast, the A_5_–P_10_ hydrogel did not exhibit this behavior until approximately 50% strain (**Figure 2c iv**). When handling these hydrogels, the first three compositions were rubbery and solid, but the A_5_–P_10_ was sponge-like and exuded much water upon deformation.

**Figure 2.**
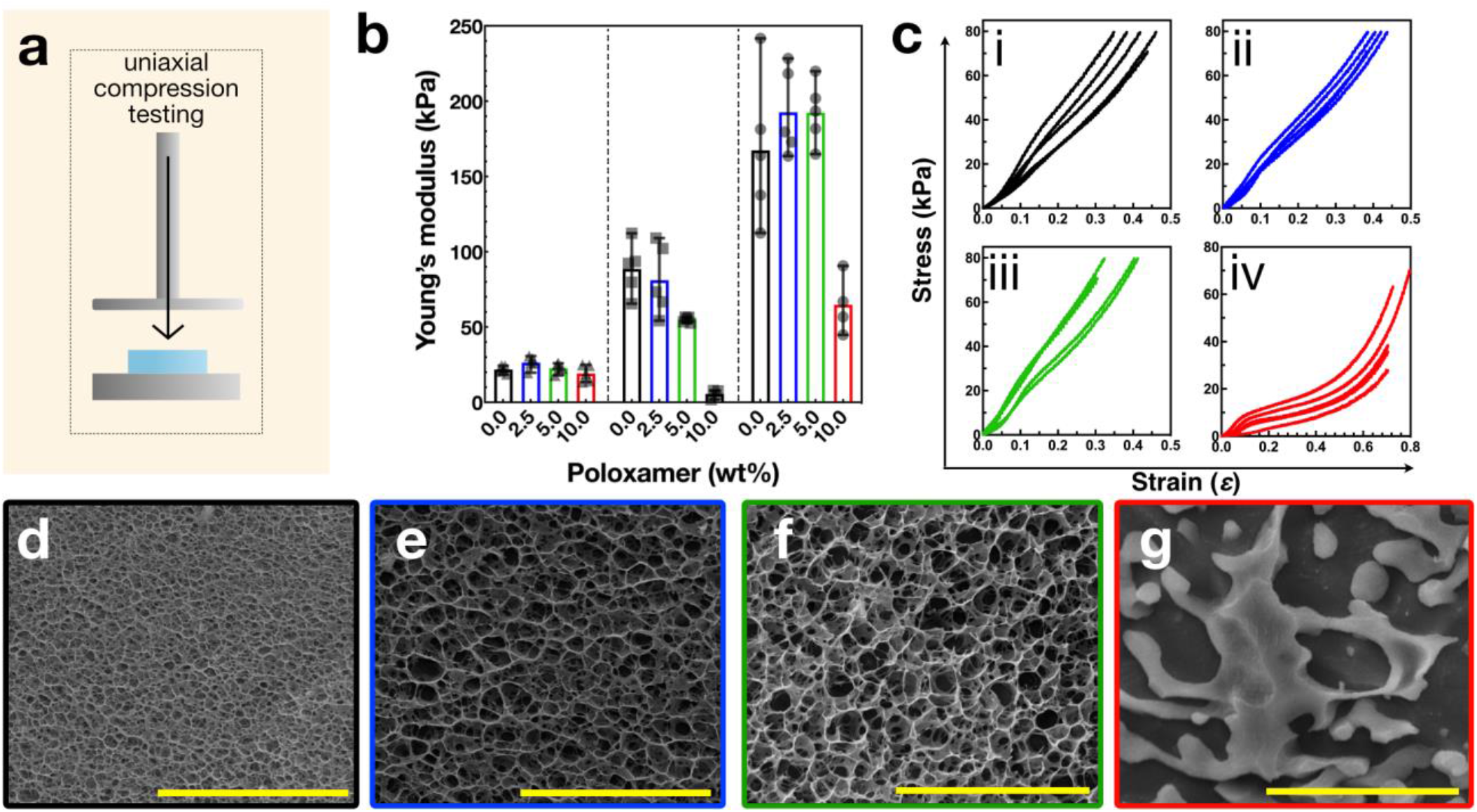
Mechanical and structural analysis of porous alginate hydrogels. **a** schematic of the unconfined uniaxial compression test. Samples were compressed at 1 mm·min^−1^. **b** Young’s moduli from compression tests of low_alg_ (triangles, left), med_alg_ (squares, middle), and high_alg_ (circles, right) hydrogels. Poloxamer concentration indicated as 0.0 (black), 2.5 (blue), 5.0 (green), and 10.0 wt% (red) bars. **c** Stress–strain curves for the high_alg_ hydrogels across all Poloxamer concentrations: i 0.0 wt%; ii 2.5 wt%; iii 5.0 wt%; and iv 10.0 wt%. Cryo-electron micrographs of fracture surfaces of the high_alg_ gels with varying Poloxamer concentrations. **d** 0 wt% Poloxamer. **e** 2.5 wt% Poloxamer. **f** 5 wt% Poloxamer. **g** 10 wt% Poloxamer. Scale bars = 10 µm.

Next, the effect of the Poloxamer porogen on hydrogel network structure was assessed in the A_5_ group by imaging fracture surfaces of flash frozen samples *via* cryo-electron microscopy, to preserve the structure of the hydrated state as much as possible. The micrographs of the A_5_–P_0_, –P_2.5_, and – P_5_ hydrogels revealed tightly organized, three-dimensional networks with pore sizes visibly enlarged in the hydrogels formed with Poloxamer (**Figure 2d i–iii**). Measurements of the pore size diameters revealed a significant increase from 0.6 ±0.2 µm to 1.1 ±0.3 and 1.3 ±0.4 µm when Poloxamer porogen was present during hydrogel fabrication (**Supplementary figure S4**), confirming that Poloxamer can be used to adjust the pore size of alginate hydrogels. In the sample containing the highest porogen concentration, A_5_–P_10_, these intricate structures were absent and instead the alginate network appeared amorphous, composed of dense regions of alginate separated by large voids. This loss of structure likely occurred due to the formation of a hierarchical pore structures due to the high porogen concentration, which prevented a continuous alginate network from forming, thus explaining the loss of mechanical strength observed in compression testing.

### Mooney–Rivlin model of hyperelasticity

The data from the compression testing was modelled to reveal it fit a two-parameters Mooney– Rivlin model of hyperelasticity (**Equation 4**)^13^:

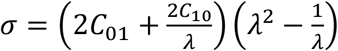

**Equation 4** Two-parameters Mooney-Rivlin hyperelastic relation between the uniaxial stress σ, the parameters C_01_ and C_10_ and the elongation λ.

The hydrogel systems show typical Drucker-type stability characteristics, with C_0_ always positive and the total of the two parameters higher than zero^14^. The compressive stress-strain curves also exhibit no inflection point (**Figure 3a**), indicating that two- or three-parameter Mooney-Rivlin models are sufficient to simulate these alginate/poloxamer systems. The relationship between the Mooney-Rivlin models and the material’s Young’s modulus is given by 6(*C*_01_+*C*_10_)^15^, which aligns with the actual Young’s moduli measured during the tests (**Figure 2**). The C_01_ parameter is particularly sensitive to alginate content, peaking at 5 wt%. The highest magnitudes of the Mooney-Rivlin constants are observed for the 5 wt% alginate and 5 wt% poloxamer combination (**Figure 3b-c**). This supports the findings from the quasi-static Young’s modulus tests, indicating that this specific composition of alginate and poloxamer enhances compressive stiffness by up to 30% at strain.

**Figure 3.**
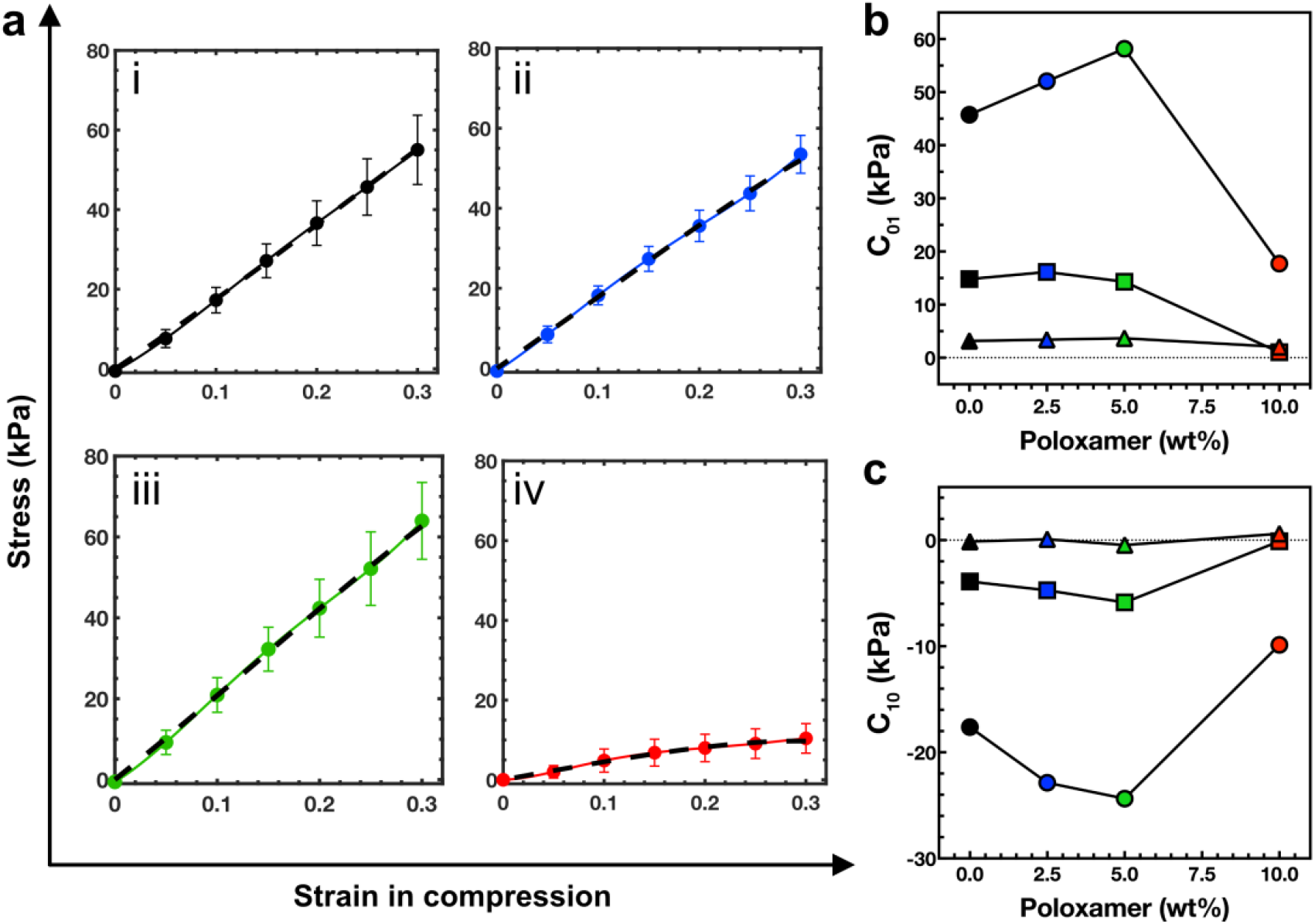
Mooney–Rivlin model of hyperelasticity. **a** The quasi-static uniaxial compression data was fitted to a Mooney–Rivlin model of hyperelasticity. The displayed curves correspond to the 5 wt% alginate hydrogels with Poloxamer at 0 wt% (i), 2.5 wt% (ii), 5 wt% (iii), and 10 wt% (iv). Solid lines with circles indicate the experimental data and the dashed lines indicate the Mooney–Rivlin model. Please refer to the supplementary information for the fits of the 1.25 wt% and 2.5 wt% alginate hydrogels. **b** The C_01_ parameter of the Mooney–Rivlin model for the 1.25 wt% alginate (triangles), 2.5 wt% alginate (squares), and 5.0 wt% alginate (circles) at increasing Poloxamer concentrations of 0 wt% (black), 2.5 wt% (blue), 5.0 wt% (green), and 10.0 wt% (red). **c** The C_10_ parameter of the Mooney–Rivlin model, as in **b**.

### Porous hydrogel with significant vibration damping properties

The A_5_ hydrogels were subjected to vibration transmissibility tests following an ISO 10819:2013 standard approach to explore their dynamic mechanical properties, determine their potential for mechanical energy dissipation, and investigate the effect of porosity on these properties. From such experiments, the dynamic modulus (*E*_*d*_) and dynamic loss factor (*η*_*d*_), can be determined. Hydrogel samples were prepared *via* the dialysis-casting method, as described above, to provide cuboid blocks with dimensions 30×30×15 mm (**Figure 4a, i–ii; Supplementary figure S3**). To perform the vibration transmissibility experiments, a hydrogel block was sandwiched between an aluminum baseplate, mounted onto an electrodynamic shaker, and a connecting plate, on to which increasing masses could be systematically installed to pre-stress the hydrogel blocks (**Figure 4a, iii**). Finally, accelerometers attached to the baseplate and top mass recorded the vibration wave amplitude as a function of acceleration across five increasing acceleration rates (1.2–5.9 m^−2^), as the hydrogel samples experienced low frequency seismic vibration at 0–1000 Hz, focusing on small amplitude deformations in a low strain range of approximately 0.3–3%. The ratio of these amplitudes was calculated to determine the transfer function (TF; **Figure 4b**)^10^, with the peak of the transfer function corresponding the resonance frequency (*ω*) of the hydrogel block. Given that the stiffness of the sample is directly related to *ω* and peak of the TF, the dynamic modulus (*E*_*d*_), loss modulus (*E*_*l*_), and loss factors (*η*) could be determined (**Equations 2–5**) and were compared across different acceleration rates. *E*_*d*_ was consistent across all accelerations (**Supplementary figure S5**), indicating the hydrogels exhibited acceptable linearity in the vibration amplitude range applied.

**Figure 4.**
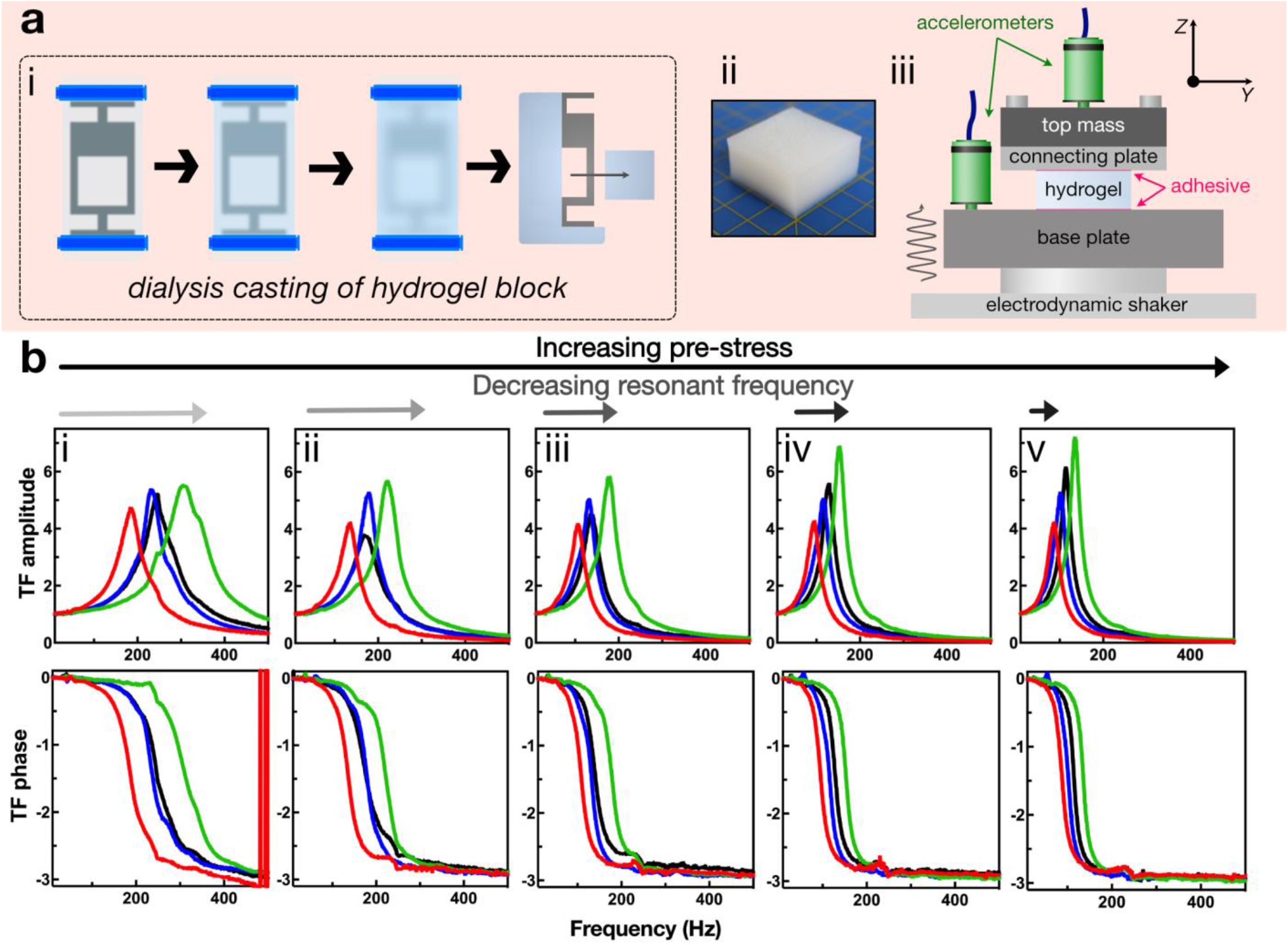
Vibration damping properties of A_5_ hydrogels **a** i: hydrogel block for vibration transmissibility experiments were prepared *via* dialysis [see Figure 1a–d]. ii: Optical image of a hydrogel block of dimensions 30×30×15 mm. Yellow grid demarcates 1 cm. iii: Schematic of the experimental set-up on an electrodynamic shaker. The hydrogel block was affixed between the base plate and connecting plate with alginate tray adhesive (magenta). A top mass could be attached to the connecting plate *via* screws, and increasingly heavy top masses could be interchanged to apply increasing pre-compressive strain to the sample prior to vibration. Accelerometers (green) were connected to the base plate and top mass to measure the amplitude of mechanical waves before and after transmission through the hydrogel. **b** Vibration transmissibility data. TOP: Representative transmissibility factor (TF) amplitude curves of 0.0 (black), 2.5 (blue), 5.0 (green), and 10.0 (red) wt% Poloxamer hydrogels. Measurements were taken under five levels of pre-compression due to increasing load with the top mass: i) 32.44 g; ii) 64.25 g; iii) 111.10 g; iv) 160.43 g; and v) 209.13 g. BOTTOM: corresponding TF phase curves.

Comparing the peaks of the transfer functions of the four A_5_ hydrogels, decreasing resonance frequency correlated with increasing pre-stress (**Figure 4b, i–v**). Compared to the A_5_–P_0_ control, A_5_– P_2.5_ exhibited negligible difference (black and blue lines, respectively), but the transfer function peaks of the A_5_–P_5_ hydrogel were of higher frequency and amplitude. For example, with a top mass of 111.10 g (**Figure 4b, iii**), the resonance frequency for the control A_5_–P_0_ hydrogel was 175 Hz and peaked at 4.5, whereas for the more porous A_5_–P_5_, the resonance frequency was 200 Hz and peaked at 6. In contrast, the transfer function of the softer A_5_–P_10_ hydrogels were peaked at lower resonance frequency and with lower magnitudes, compared to the other hydrogels (**Figure 4b, i–v** red line). Accordingly, the architecture of the hydrogel network is important. Given that resonance frequency is directly related to stiffness (**Equation 3**), these data suggest that the increased porosity of the A_5_– P_5_ had the effect of pushing the alginate into a smaller volume fraction and may have reinforced and stiffened the alginate network, although it should be noted that the A_5_–P_2.5_ hydrogel did fit this trend.

In each of the A_5_ hydrogels, *E*_*d*_ was observed to trend upwards with increasing pre-stress (and decreasing resonant frequency), likely due to higher dynamic loading (**Figure 5a; Supplementary figure S6**). Importantly, the dynamic moduli were found to be approximately >10–20-fold greater than the corresponding Young’s moduli. For example, in the A_5_–P_0_ hydrogel, under the lowest degree of pre-stress, *E*_*d*_ was 1.6 MPa, whereas the Young’s modulus measured in quasi-static compression testing was 160 kPa. As the degree of pre-stress increased, so too did *E*_*d*_, up to ∼2 MPa at 3% dynamic strain. Similar trends were observed for all the hydrogels. The stiffer A_5_–P_5_ hydrogel exhibited an initial *E*_*d*_ of 2.5 MPa at resonance with low levels of pre-stress, which increased to 2.8 MPa. The soft, structureless A_5_–P_10_ hydrogel was also substantially stiffer at resonance, exhibiting an *E*_*d*_ of 0.9 MPa that increased to 1.1 MPa under compression. The loss of the repetitive network structure led to a decrease in dynamic modulus (**Figure 5b**). The A_5_–P_2.5_ hydrogel did not however exhibit a dynamic modulus with an intermediate value between the A_5_–P_0_ and A_5_–P_5_ samples, but instead was lower. We postulate that the underlying reason for this phenomenon may be associated with the critical micelle concentration of Poloxamer 407, which is close to 4 wt%^16^. This implies that a complete phase transition may not have occurred, leading to a potential interaction with the alginate that could influence the cross-linking density. Alternatively, the cryo-EM micrographs reveal a more closed cell architecture for the A_5_–P_0_ hydrogel, whereas the A_5_–P_5_ hydrogel exhibits a more open structure. The A_5_–P_2.5_ composition closely resembles the control; however, the openness of the A_5_– P_5_ structure is more pronounced, overshadowing this similarity.

**Figure 5.**
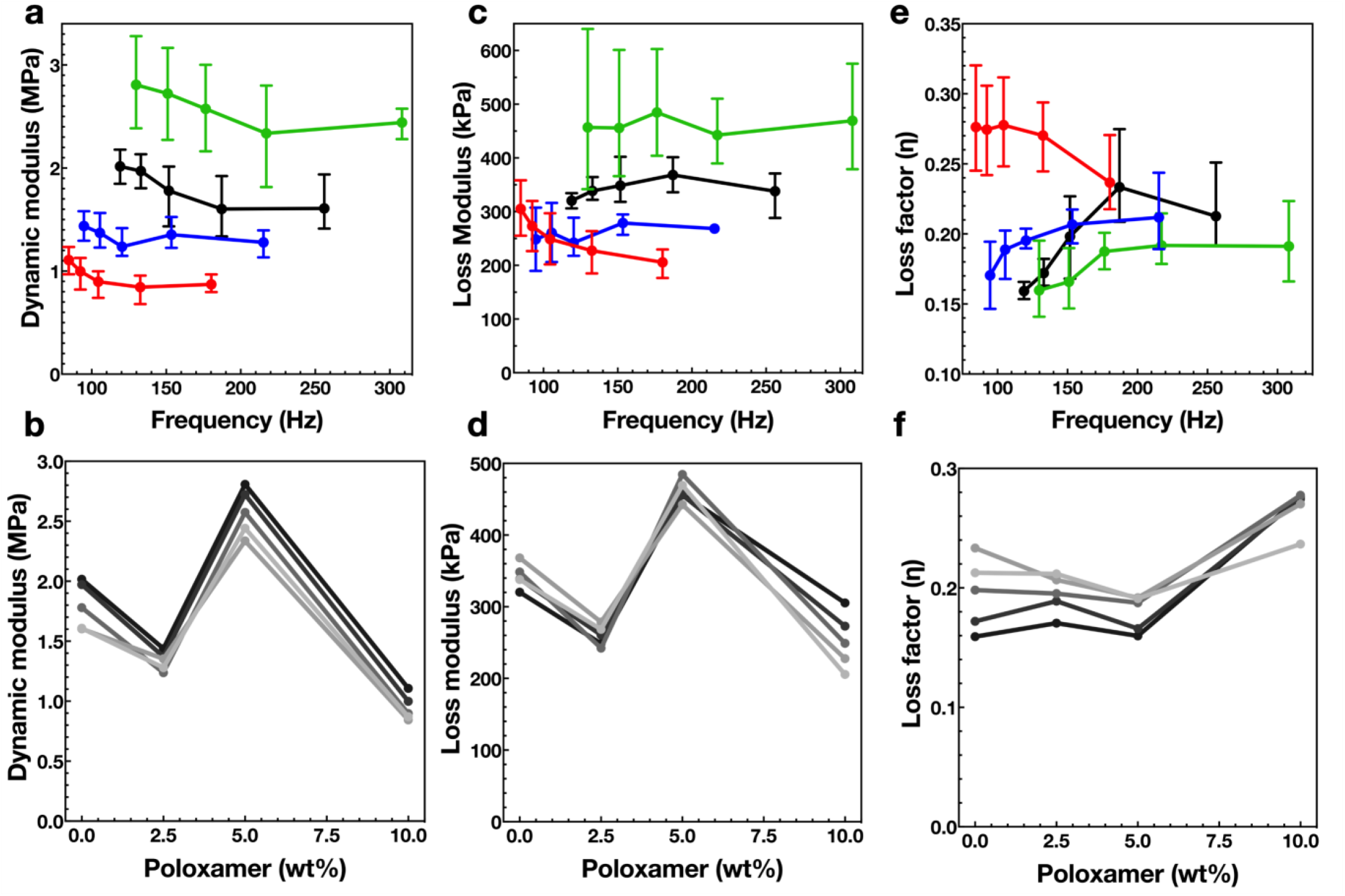
Vibration damping properties of the A_5_ hydrogels. **a** Dynamic modulus of the A_5_–P_0_ (black), A_5_–P_2.5_ (blue), A_5_– P_5_ (green), and A_5_–P_10_ (red) Poloxamer hydrogels, plotted at resonant frequency. **b** The corresponding loss modulus of the A_5_ hydrogels. **c** Calculated loss factors (*η*) for the A_5_ hydrogels. **d** The dynamic modulus plotted against porogen weight %, under increasing pre-strain corresponding to increasing top masses, from light to dark grey. **e** The loss modulus plotted against porogen weight %, under increasing pre-strain corresponding to increasing top masses, from light to dark grey. **f** Loss factors (*η*) plotted against porogen weight %, under increasing pre-strain corresponding to increasing top masses, from light to dark grey. Error bars in **a–c** correspond to range, n = 3.

The loss modulus from the vibration transmissibility tests trended downwards, with increasing pre-compression in the A_5_–P_0_ hydrogel from a peak of 400 kPa to 320 kPa (**Figure 5c**). Whilst the loss moduli of the stiff A_5_–P_5_ hydrogel was linear, the soft A_5_–P_10_ sample increased from 200 to 300 kPa. Consequently, the loss factor (*η*), the ratio of loss modulus to dynamic elastic modulus describing the proportion of the mechanical energy dissipated, exhibited different trends. In the control alginate-only hydrogel, the loss factor decreased steeply at higher pre-compression from 0.24 to 0.16. The trend of the A_5_–P_2.5_ hydrogel was similar. The stiffer A_5_–P_5_ hydrogel exhibited lower loss factors which plateaued at low pre-compression at 0.19 and decreased to 0.16 at higher pre-compression. Interestingly, the loss factor of the A_5_–P_10_ hydrogel was higher and increased from 0.24 to a plateau of 0.28 at higher pre-compression.

DMA tests on the A_5_-P_5_ hydrogels, conducted in the 100 Hz to 130 Hz frequency range, reveal storage modulus values between 2.3 MPa and 2.5 MPa, with tan δ values from 0.15 to 0.18 (**Supplementary figure S7**). These findings align with the vibration transmissibility tests, confirming a high loss modulus in these hydrogel systems within the examined frequency ranges.

The high dynamic moduli and loss factors observed in these porous hydrogels were in part due to the pneumatic/hydrostatic forces exerted through the porous network, similarly to the dissipative and strain-rate effects observed in polymeric closed-cell foams with entrapped gas^17^. The dynamic stiffening effect in these gels is remarkable. For comparison, open-cell polyurethane foams with similar static Young’s moduli (120-150 kPa) show only a 50% increase in dynamic modulus during vibration transmissibility tests^10^. These hydrogels will exhibit the ability to dampen through several mechanisms, including the interaction between solvent and polymer fibers, and polymer–polymer friction.

Alginates are also known to exhibit viscoelastic properties and can dissipate energy through breaking and reforming of the ionic cross-links holding the network together, permitting network remodeling under stress^18^. Comparing the A_5_–P_0_ and A_5_–P_5_ hydrogels, increasing the size of the macropores in the alginate network resulted in reinforcing the network fibers and increased their effecting cross-linking density. This reinforcement resulted in a stiffer network that resisted dynamic deformation, as observed in the increased dynamic moduli.

## Conclusions

The alginate/poloxamer systems developed in this work exhibit multiscale porosity, enabling poroelastic and enhanced damping effects in the 50 Hz to 300 Hz frequency range. These frequencies correspond to low-frequency vibrations commonly encountered in machinery and transport systems, which are typically challenging to dampen using conventional elastomeric and polymeric materials^5^. The dynamic stiffening of these gels within the considered frequency range is notable, with increases of more than an order of magnitude compared to their quasi-static Young’s modulus. This significant rise in dynamic modulus, observed using a vibration transmissibility rig, is further validated by cyclic compression tests conducted with a dynamic mechanical analyser. The alginate/poloxamer system is tunable, with both mechanical and dynamic properties sensitive to the weight fractions of poloxamer and alginate. The average loss factors range from 16% to 26%. A weight fraction of 5% for both alginate and poloxamer yields the stiffest hydrogels, with quasi-static moduli around 180 kPa and dynamic moduli up to 2.8 MPa. The loss factors for the A5-P5 gels range from 16% to 19%, while higher loss factors of up to 28% can be achieved with different weight fractions, although this results in lower quasi-static and dynamic moduli.

While the hydrogels developed here are produced via dialysis, the alginate/poloxamer systems can also be bioprinted^9^, providing flexible manufacturing and deposition options with additive techniques. Additionally, these hydrogels are made from biosourced and biodegradable materials, offering a sustainable alternative to conventional fossil-based damping materials on the market.

## Acknowledgements

This work has been funded by the UK Defense Science and Technology Laboratory (Grant ID: DSTL0000024791), the Office of Naval Research Global (Grant ID: N62909-201-2061), and the ERC H2020 AdG NEUROMETA project (01020715). The authors would also like to express their gratitude to Professor Petra C. Oyston, Mr Matthew D Eagling (Dstl), and Drs Patrick P. Rose and Scott A. Walper (ONR G) for their support and insightful discussions throughout the project. The authors thank Judith Mantell of the Wolfson Bioimaging Facility for their assistance in capturing the cryo-EM micrographs presented in this work.

## Notes

### Competing Interest Statement

The authors have declared no competing interest.

