## Supplementary information for "Tunable porosity in a hydrogel with extreme vibration damping properties"

### Supplementary figures

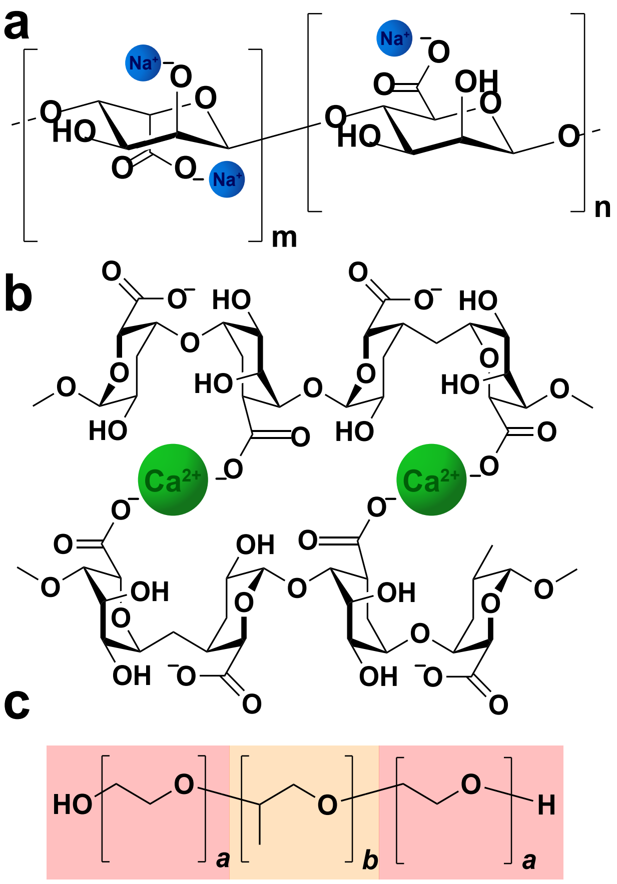


**Supplementary figure S1**. Chemical structures of the polymers used to make porous hydrogels. **a** Linear alginate polymer comprised of (1→4)-linked β-D-mannuronate (m) and α-L-guluronate (n), bound to monovalent sodium ions ($Na^+$; blue spheres). **b** Guluronate chains of alginate cross-linked by divalent calcium ions ($Ca^{2+}$; green spheres). **c** Structure of the triblock copolymer Poloxamer 407, comprised of hydrophilic polyethylene glycol (red boxes) and hydrophobic propylene glycol (orange box). *a* = 101 and *b* = 56.

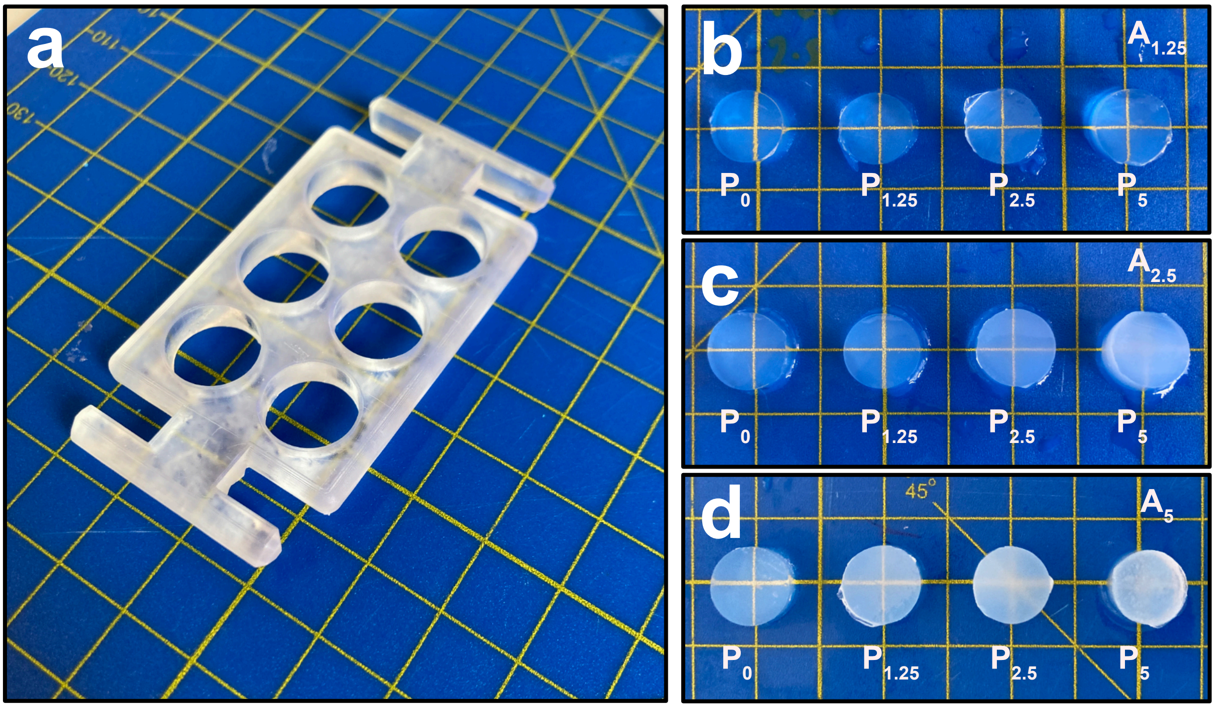


**Supplementary figure S2.** Forming hydrogels for compression testing. **a** The mold used in dialysis to yield hydrogel discs 12 mm in diameter and 5 mm in height. **b** Optical images of the $A_{1.25}$ hydrogel discs with increasing Poloxamer concentration. **c–d** Corresponding images of the $A_{2.5}$ and $A_5$ hydrogels, respectively. Yellow lines demarcate 1 cm.

---

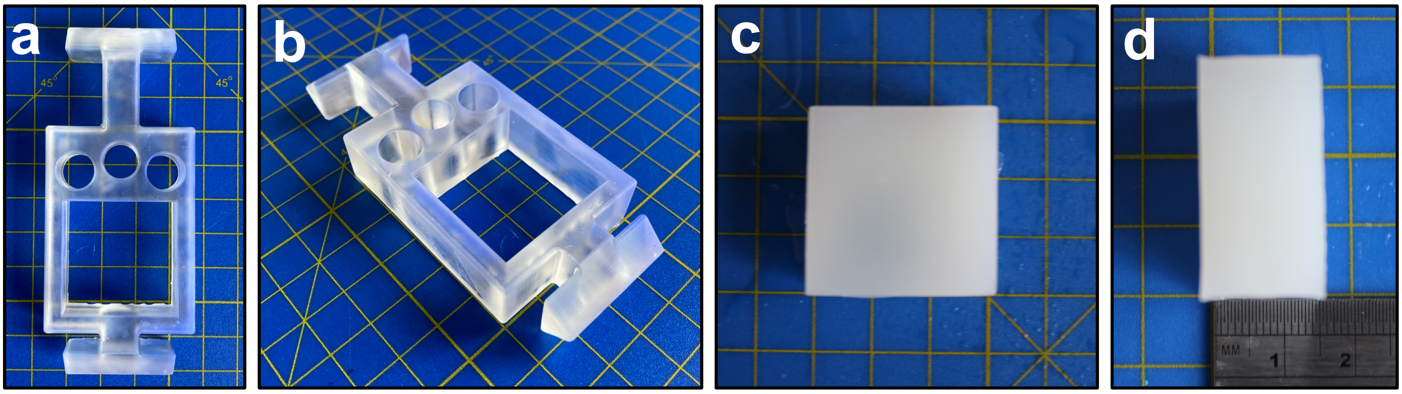


**Supplementary figure S3.** Forming the hydrogels for dynamic mechanical characterisation. **a–b** Optical images of the mold used in dialysis to form hydrogel blocks 30×30×15 mm. **c–d** Images of the blocks.

---

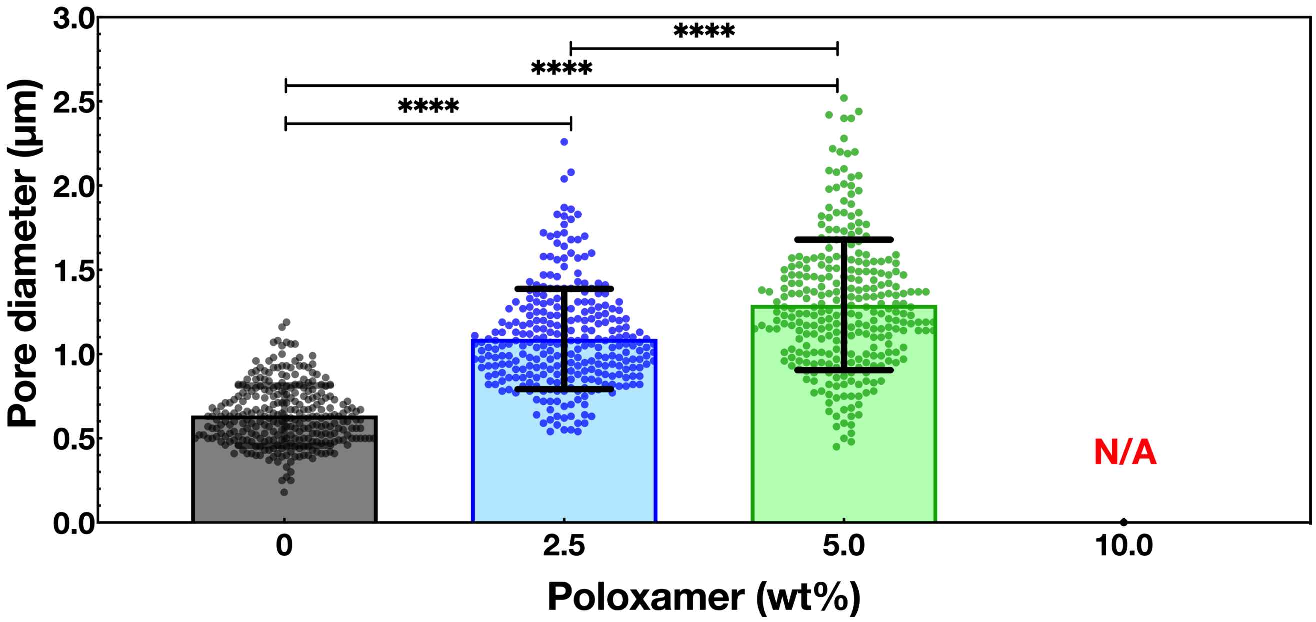


**Supplementary figure S4.** Pore diameters of the $A_5$ hydrogels. Average pore diameter and distribution measured from the cryo-EM micrographs. n = 290, error bars = SD. **** = p <0.0001.

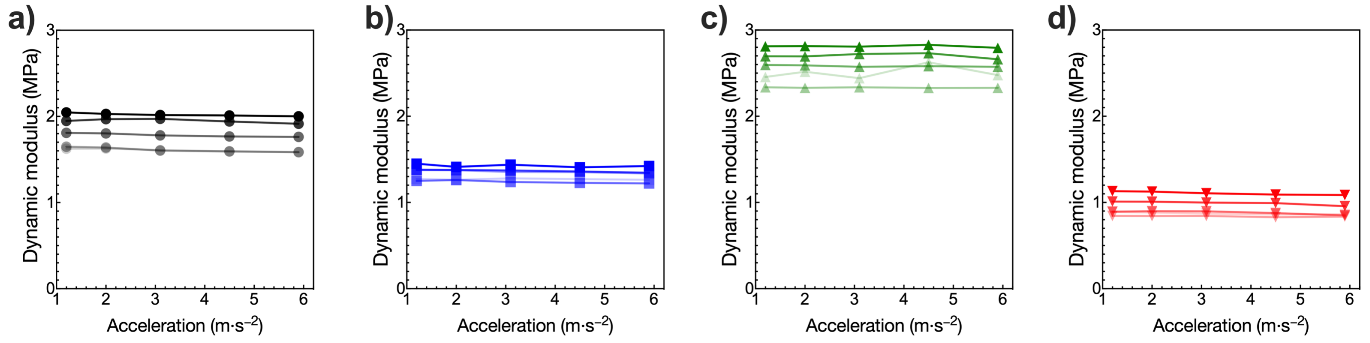


**Supplementary figure S5.** Linearity of the dynamic mechanical analysis set-up. The dynamic modulus of $A_5$–$P_0$ (**a**; black), $A_5$–$P_{2.5}$ (**b**;blue), $A_5$–$P_5$ (**c**;green), and $A_5$–$P_{10}$ (**d**;red) at increasing acceleration rates under increasing degrees of pre-stress (increasing opacity).

---

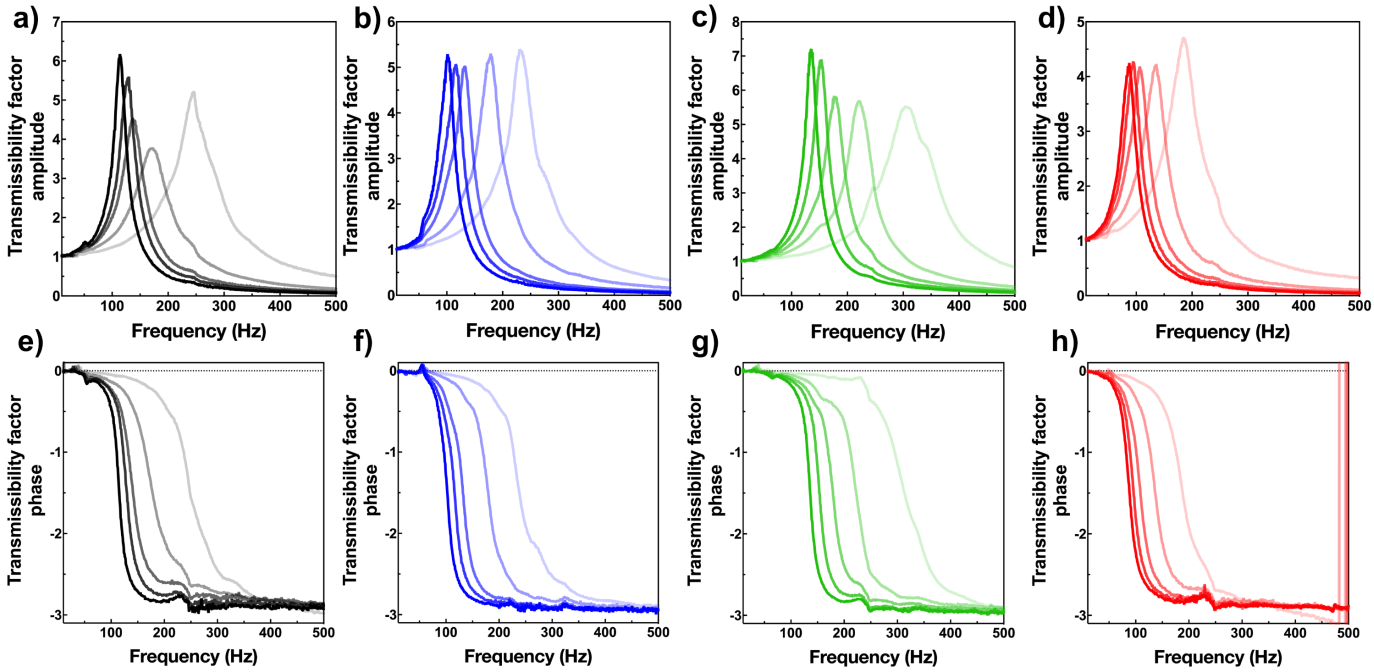


**Supplementary figure S6.** Transmissibility factor amplitudes (**a–d**) and corresponding phase curves (**e–h**) of $A_5$–$P_0$ (black), $A_5$–$P_{2.5}$ (blue), $A_5$–$P_5$ (green), and $A_5$–$P_{10}$ (red). Increasing opacity corresponds to increasing pre-stress. This data is also displayed in Figure 5b, but re-plotted here per individual hydrogel composition.

---

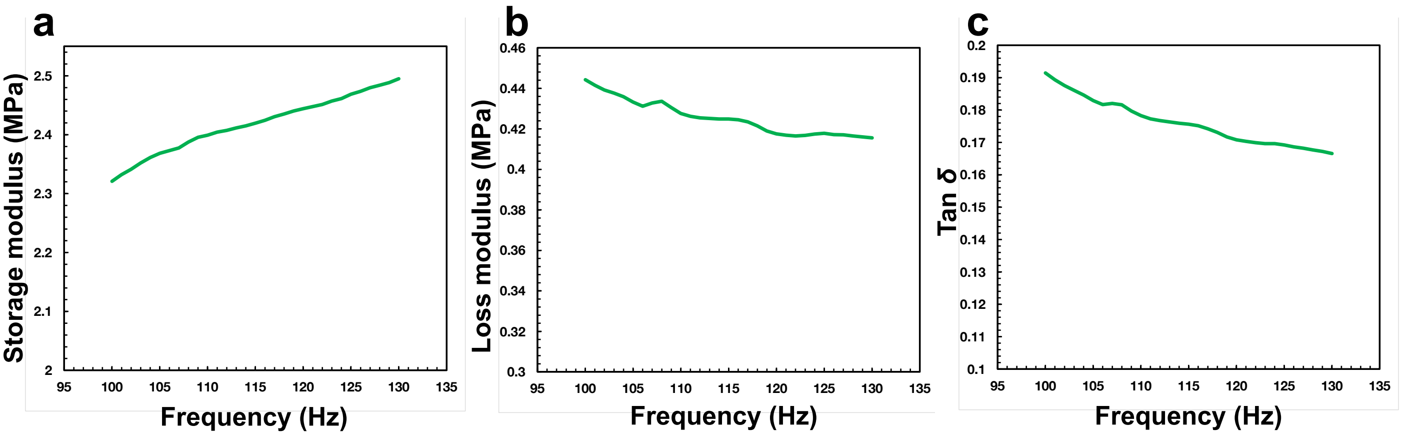


**Supplementary figure S7.** Characterization of $A_5$–$P_5$ hydrogel using a dynamic mechanical analyzer (DMA) between 100–130 Hz. **a** Storage modulus. **b** Loss modulus. **c** Tan $\delta$. Tests were done at 25 °C, with a pre-loading force of 0.001 N and oscillation amplitude of 10 μm.

### Supplementary tables

**Supplementary table S1.** The mass of alginate, Poloxamer 407, and water required to make 50 g gels for dialysis casting.

| *Hydrogel* | *Alginate mass (g)* | *Poloxamer 407 mass (g)* | *Water mass (g)* |
|---|---|---|---|
| **$A_{1.25}$–$P_0$** | 0.625 | 0.0 | 49.375 |
| **$A_{1.25}$–$P_{2.5}$** | | 1.25 | 48.125 |
| **$A_{1.25}$–$P_5$** | | 2.5 | 46.875 |
| **$A_{1.25}$–$P_{10}$** | | 5.0 | 44.375 |
| **$A_{2.5}$–$P_0$** | 1.25 | 0.0 | 48.75 |
| **$A_{2.5}$–$P_{2.5}$** | | 1.25 | 47.5 |
| **$A_{2.5}$–$P_5$** | | 2.5 | 46.25 |
| **$A_{2.5}$–$P_{10}$** | | 5.0 | 43.75 |
| **$A_5$–$P_0$** | 2.5 | 0.0 | 47.5 |
| **$A_5$–$P_{2.5}$** | | 1.25 | 46.25 |
| **$A_5$–$P_5$** | | 2.5 | 45.0 |
| **$A_5$–$P_{10}$** | | 5.0 | 42.5 |

**Supplementary table S2.** Derived coefficients from the Mooney–Rivlin model of hyperelasticity.

| *Alginate (wt%)* | *Poloxamer (wt%)* | *$C_{01}$ (kPa)* | *$C_{10}$ (kPa)* |
|---|---|---|---|
| **1.25** | 0.0 | 3.17 | -0.13 |
| | 2.5 | 3.41 | 0.08 |
| | 5.0 | 3.67 | -0.47 |
| | 10.0 | 2.03 | 0.60 |
| **2.5** | 0.0 | 14.81 | -3.90 |
| | 2.5 | 16.15 | -4.72 |
| | 5.0 | 14.31 | -5.87 |
| | 10.0 | 1.02 | -0.11 |
| **5.0** | 0.0 | 45.75 | -17.61 |
| | 2.5 | 52.06 | -22.87 |
| | 5.0 | 58.18 | -24.36 |
| | 10.0 | 17.72 | -9.87 |
